# Mismatch-repair signature mutations activate gene enhancers across colorectal cancer epigenomes

**DOI:** 10.1101/411264

**Authors:** Stevephen Hung, Alina Saiakhova, Zachary Faber, Cynthia F. Bartels, Devin Neu, Ian Bayles, Evelyn Ojo, Ellen S. Hong, W. Dean Pontius, Andrew R. Morton, Ruifu Liu, Matthew Kalady, David N. Wald, Sanford Markowitz, Peter C. Scacheri

## Abstract

Commonly-mutated genes have been found for many cancers, but less is known about mutations in *cis*-regulatory elements. We leverage gains in tumor-specific enhancer activity, coupled with allele-biased mutation detection from H3K27ac ChIP-seq data, to pinpoint potential enhancer-activating mutations in colorectal cancer (CRC). Analysis of a genetically-diverse cohort of CRC specimens revealed that microsatellite instable (MSI) samples have a high indel rate within active enhancers. Enhancers with indels show evidence of positive selection, increased target gene expression, and a subset is highly recurrent. The indels affect short homopolymer tracts of A/T and increase affinity for FOX transcription factors. We further demonstrate that signature mismatch-repair (MMR) mutations activate enhancers using a xenograft tumor metastasis model, where mutations are induced naturally via CRISPR/Cas9 inactivation of *MLH1* prior to tumor cell injection. Our results suggest that MMR signature mutations activate or augment enhancers in CRC tumor epigenomes to provide a selective advantage.

## Introduction

In the past decade, tumor sequencing efforts by TCGA, ICGC, and others have identified the most frequently mutated driver genes in most common forms of cancer (Lawrence et al. 2014; Kandoth et al. 2013). Outside protein-coding genes, mutations in non-coding regions that disrupt enhancer-gene control are emerging as a major mechanism of cancer development. Examples include large chromosomal rearrangements that hijack enhancers to oncogenes, such as the Burkitt lymphoma translocation that repositions IgH enhancers upstream of *MYC* (Battey et al. 1983). Copy number alterations can amplify enhancer sequences near oncogenes. Deletions can remove boundaries between enhancers and proto-oncogenes, and inversions can flip enhancers to proto-oncogenes (Zhang et al. 2016; Beroukhim et al. 2016; Hnisz et al. 2016). Besides large structural variants that rewire gene-enhancer interactions, small-scale mutations that lie *within* regulatory elements and alter their activity can occur. The first of these to be discovered were recurrent point mutations in the *TERT* promoter in melanoma and other cancers (Huang et al. 2013). Other examples include a small indel that creates a super-enhancer that drives the overexpression of the *TAL1* oncogene in T-ALL (Mansour et al. 2014), and recurrent enhancer substitutions and indels that affect the expression of *PAX5* in CLL (Puente et al. 2015). The discovery of these driver events has motivated searches for additional enhancer mutations in other common cancers, but so far their prevalence and relevance to the cancer phenotype remain largely undetermined.

The identification of functional enhancer mutations is challenging due to several confounding factors. First, mutation rates vary considerably between different tumor types and even among tumors of the same subtype. Second, tumor epigenomes are heterogeneous and mutation rates are profoundly influenced by chromatin states, with euchromatic early-replicating regions showing a low mutation rate relative to heterochromatic late-replicating regions (Schuster-Böckler and Lehner 2012; Polak et al. 2015). Given this variation, the conventional approach of overlaying mutations detected through tumor sequencing with a “reference” epigenome is suboptimal. Strategies that facilitate simultaneous capture of both sequence content and regulatory activity are more suitable. Third and perhaps most importantly, for most cancers the cell type of origin is unknown or unavailable for epigenomic studies. The lack of the normal comparator makes it difficult to assess whether a putative mutation influenced the activity of the regulatory element relative to the normal cell from which the tumor was derived.

Through ChIP-seq analysis of enhancer histone marks (H3K4me1 and H3K27ac), we previously compared the enhancer epigenomes of a genetically-diverse cohort of human CRC models to normal colonic crypts, the cell type of origin for CRC. We identified Variant Enhancer Loci (VELs) as sites that differed in the levels of H3K4me1 and H3K27ac between normal crypts and each CRC sample (Akhtar-Zaidi et al. 2012; Cohen et al. 2017). Here, we pinpoint functional enhancer mutations in VELs directly from H3K27ac ChIP-seq data, using the premise that a DNA variant in an enhancer with higher H3K27ac levels in CRC than normal colon cells may have contributed to the activation of that “gained” enhancer. Our analysis shows that CRC samples with underlying deficiencies in mismatch-repair harbor an exceptionally high indel rate in gained enhancers compared to their already high background mutation rate. We provide evidence that these non-coding mutations, previously presumed to be inert, are functional.

### Identification of putative enhancer activating indels

We looked for candidate mutations that augment enhancer activity by identifying somatic mutations in regions with elevated levels of H3K27ac in CRC relative to normal colon (Figure 1a). A key step in the analysis is identifying instances of allele bias, where H3K27ac ChIP-seq read depth is higher on the allele containing the mutation than on the reference allele. We further eliminate mutations that are not predictive of gained H3K27ac enrichment (i.e., the mutation occurs in a cell line with the gained enhancer, but not in other cell lines with that same enhancer), as these are more likely to represent passenger events. We focused on indels as these have previously been shown to stimulate enhancer activation through *de novo* creation of transcription factor binding sites (Mansour et al. 2014). Using H3K27ac ChIP-seq data from 24 cell lines derived from all clinical stages of CRC, we detected a total of 355 candidate enhancer-activating indels (example shown in Figure 1b). Other signature features of enhancers were also often present at these sites. Specifically, 85% (301/355) were located distal (>5 kb) to transcription start sites (Figure 1c), 76% overlapped H3K4me1 peaks, and 73% were located <1 kb from a DNaseI hypersensitive site. Nineteen out of 20 indels detected through our analysis pipeline were validated through Sanger sequencing, indicating high specificity of our method (Figure 1-figure supplement 1a-b).

**Figure 1:**
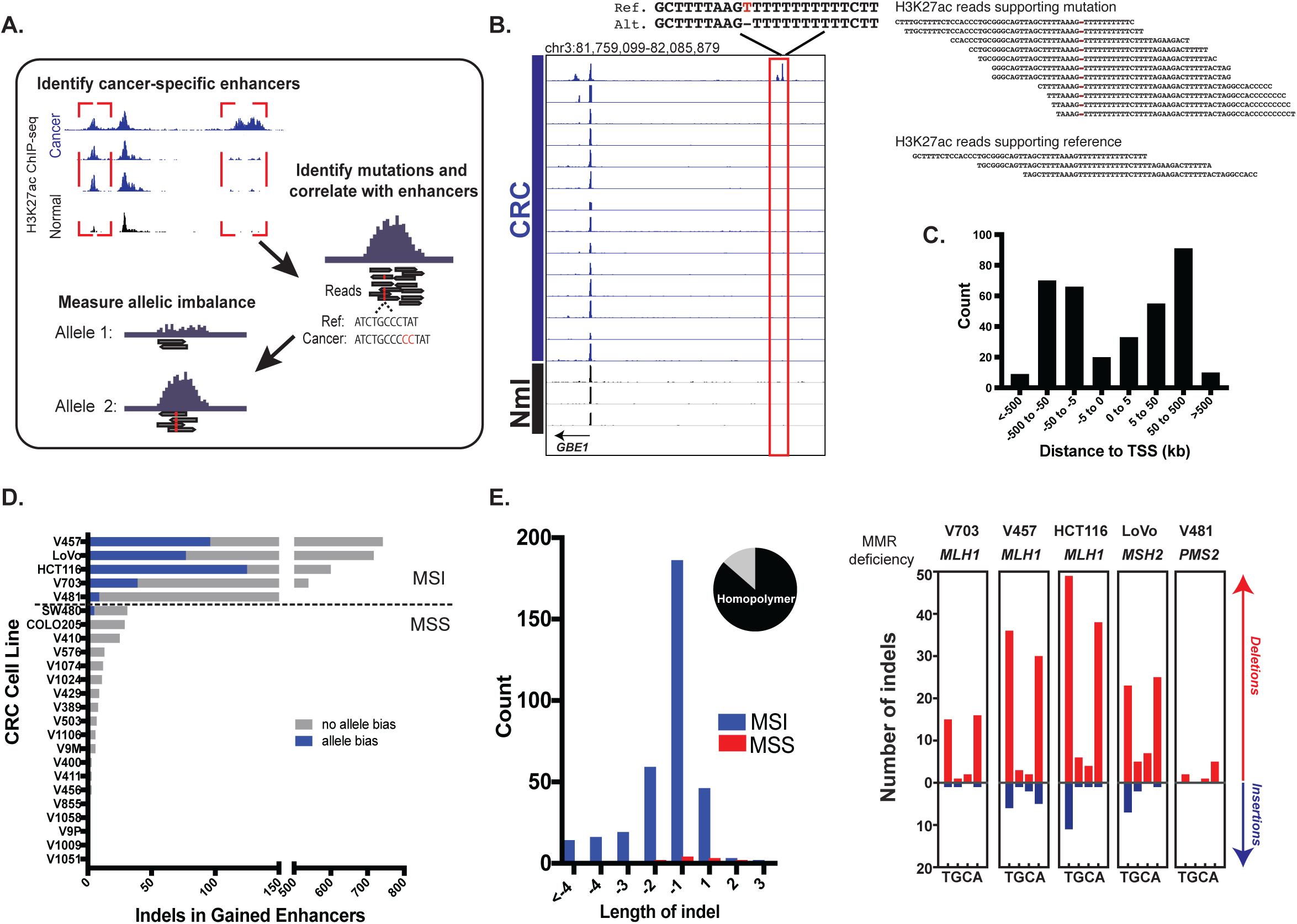
Identification of functional enhancer mutations. **A.** Overview of approach. **B.** Genome Browser snapshot of a putative enhancer-activating indel detected at a CRC-specific H3K27ac peak. Y-axis scales are all 0 to 25. Locus shown corresponds to chr3:81,759,099-82,085,879 (hg19). **C.** Distribution of CRC-specific H3K27ac peaks containing indels relative to transcription start sites. **D.** Number of gained enhancer-associated indels with and without allele bias detected in each CRC cell line. List of enhancer mutations is provided in Figure 1-source data 1. **E.** (left) Distribution of indel lengths detected in gained enhancers in MSI and MSS CRC samples. The pie chart shows the fraction of indels (304/355) detected in homopolymers. (right) Number of insertion-(blue) and deletion-(red) mutations in homopolymers of T, G, C, or A in the five MSI lines.

An analysis of the distribution of enhancer indel mutations across the full CRC cohort revealed two main classes, stratified by microsatellite stability (Figure 1d, Figure 1-source data 1). Microsatellite stable (MSS) samples harbored a lower enhancer mutation rate than MSI samples, which are deficient in mismatch repair (MMR) genes: *MLH1, MSH2, and PMS2*. The enhancer indels in MSI samples were predominantly short (1-2 bp) contractions of homopolymer runs of T’s and A’s (Figure 1e), the classic mutational signature found in coding regions of MSI tumors (Ionov et al. 1993; Kim et al. 2013). As expected, *MLH1* and *MSH2* deficient cell lines also showed higher indel rates than *PMS2* deficient cells (Baross-Francis et al. 2001; Hegan et al. 2006). To further test the relevance of these findings, we analyzed H3K27ac ChIP-seq data from 4 primary tumors (Cohen et al. 2017) of unknown microsatellite status. One of the 4 samples, which was subsequently identified as the only MSI sample in the group, showed an elevated enhancer mutation rate and the signature MSI mutation of poly A/T homopolymers (Figure 1-figure supplement 1c). The results indicate that enhancer mutations are prevalent in MSI-forms of CRC, and they show the same MMR signature as coding regions. The presence of enhancer mutations in both cell lines and primary tumors rules out an *in vitro*-specific mechanism of enhancer activation.

### MSI enhancer mutations show evidence of positive selection

Similar to previous studies correlating point mutation rates with active histones (Makova and Hardison 2015; Kim et al. 2013), indel mutation rate and H3K27ac levels were anti-correlated in MSS CRC. In contrast, there was no overall correlation between indel rate and H3K27ac levels in the MSI samples (Figure 2a), with both H3K27ac-enriched and depleted regions showing a mutation rate nearly 100 times higher than in MSS CRC. This result is consistent with previous studies indicating that open regions of chromatin are no longer protected upon loss of MMR (Supek and Lehner 2015). This raised the question as to whether gained enhancers are a target for mutation simply because they lie in open chromatin, or if the mutations are truly functional. We reasoned that if an indel activated the enhancer, then gained enhancers would show a higher proportion of allele-biased indels compared to enhancers that are already open, i.e, enhancers shared between CRC and normal crypt. In all 5 MSI samples, the proportion of allele-biased gained enhancers was higher than that of shared enhancers (Figure 2b). Even after controlling for enhancer length and H3K27ac signal intensity, gained enhancers were more likely to contain allele-biased indels than shared enhancers (Figure 2c, Figure 2 – source data 1), indicating positive selection of the gained enhancers with indels.

**Figure 2:**
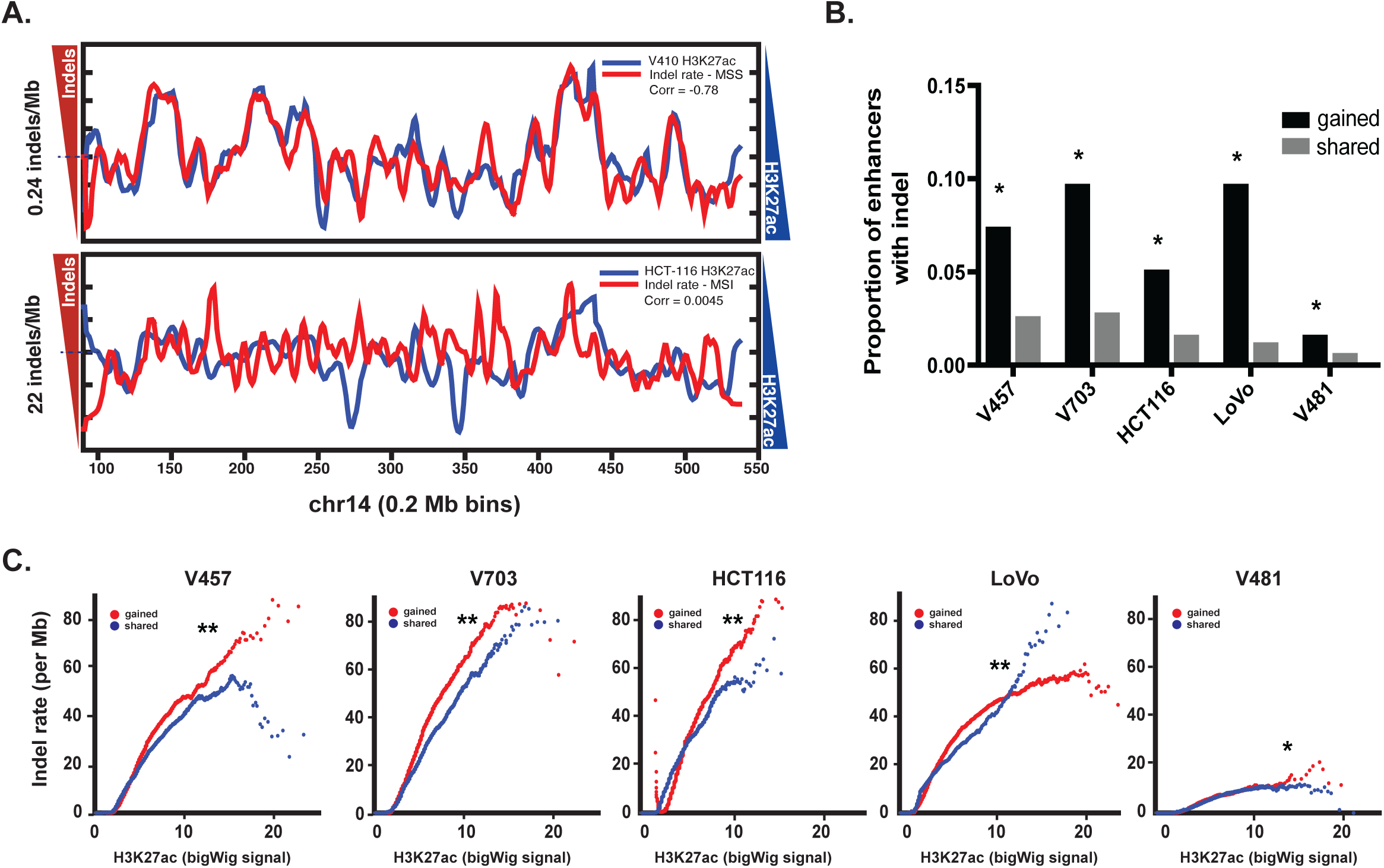
Enrichment of MSI indels in gained enhancers. **A.** MSS mutations (top) and MSI mutations (bottom) are overlaid on H3K27ac profiles (smoothed) of representative MSS (V410) and MSI (HCT-116) lines in 0.2 Mb bins on chromosome 14. Y label shows median mutation rate. The H3K27ac signal is inverted to better show anti-correlation. Pearson correlation values are shown. **B.** Proportion of gained enhancers (black) and enhancers shared with normal crypt (grey) with at least 1 allele-biased indel. * p< 1e-10, Z-test for 2 proportions. **C.** Rate of allele-biased indels (smoothed) in gained enhancers (red) and shared enhancers (blue), as a function of H3K27ac bigWig signal. **, p< 1e-10; * p< 0.05, t-test. (Figure 2-source data 1).

### MSI enhancer mutations increase expression of cancer-related genes

We next tested the effect of the indels in gained enhancers on target gene expression. To isolate the effect of the indel from that of the gained enhancer, we retrieved genes associated with gained enhancer indels, and compared their expression in cell lines with both the indel and the gained enhancer, to that of cell lines with only the gained enhancer. Genes regulated by gained enhancers containing indels were more highly expressed than those same genes regulated by gained enhancers containing the wildtype sequence (Figure 3a, Figure 3-source data 1). We further verified that this expression difference was unlikely to be due to differences in the levels of H3K27ac between the two test groups (Figure 3-figure supplement 1a). In further support of their functional relevance, genes with enhancer mutations were significantly overexpressed in primary tumors (Figure 3b). The overexpressed genes had an over-representation of genes designated as “cancer-related” by the COSMIC database, or belonging to known cancer pathways (Figure 3c). These genes include *BMP4*, a ligand of TGFB signaling often overexpressed in CRC (Duerr et al. 2012); *SKP2*, a member of the ubiquitin ligase complex implicated in lymphoma formation (Latres et al. 2001); and *SOX9*, a recurrently-mutated transcription factor in CRC that affects Wnt signaling <u>(Cancer Genome Atlas Network 2012)</u>. Gene ontology analysis revealed that amongst the top enriched functions of the indel-associated genes are cell proliferation, tissue development and embryogenesis, regulation of biosynthesis, and cell-cell signaling (Figure 3-figure supplement 1b). Collectively, the results suggest that the MSI enhancer indels lead to increased expression of target genes, several of which normally function in cell growth and early development and may therefore enhance CRC cell fitness or instigate a cell state change.

**Figure 3:**
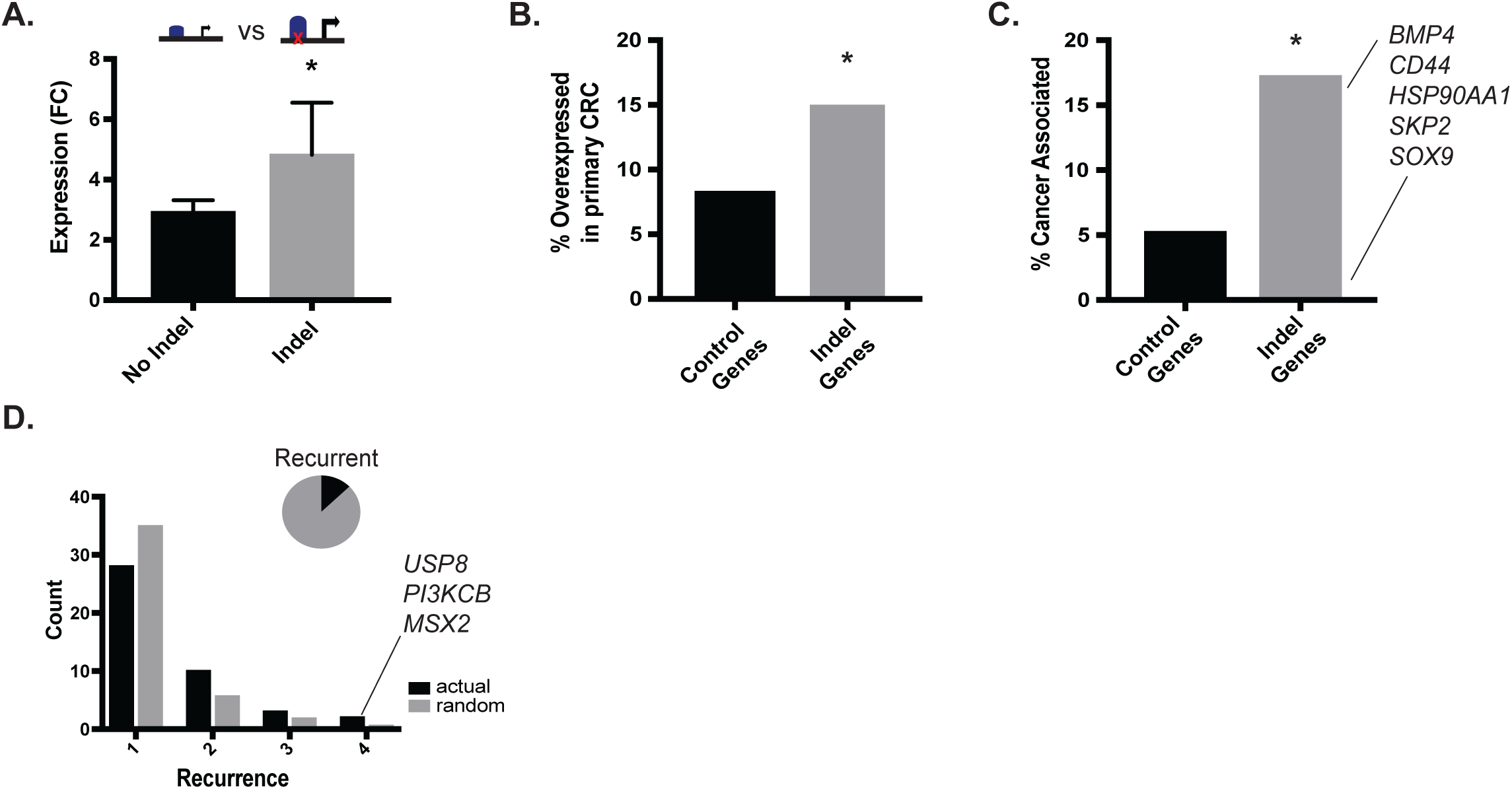
MSI enhancer indels impact target gene expression. **A.** Mean fold-change of expression (CRC/crypts) of genes associated with gained enhancers containing indels (grey; n = 484 sample-gene pairs) versus the same genes in cell lines with gained enhancers but lacking indels (black; n = 3,867 sample-gene pairs). * p< 0.01, 2-sample t-test (Figure 3-source data 1). **B.** (grey) Percent of gained enhancer indel genes in MSI cell lines that are overexpressed two-fold or more in primary CRC tumors relative to normal colon. Control (black) corresponds to the percentage of all over-expressed genes in MSI cell lines that are over-expressed two-fold or more in primary CRC. * p<0.01, Fisher exact test. **C.** (grey) Percent of over-expressed indel gained enhancer genes that have cancer-related annotations. Control (black) is the background percentage of all over-expressed genes in MSI CRC lines and primary CRC with cancer-related annotations. * p<0.05, Fisher exact test. **D.** (black) Distribution of number of samples in TCGA COAD cohort with recurrent indel. Control (grey) is average recurrence distribution from random sampling of TCGA indels (p<0.05, chi-squared test). Gene names correspond to genes whose expression correlates with enhancer signal gain. Pie chart shows the fraction of enhancer indels (46/355, or 13%) that are recurrent in at least 1 primary CRC tumor (Figure 3-source data 2).

### MSI enhancer mutations are recurrent

To test if any of the enhancer mutations were recurrent, we analyzed whole genome sequence data from 47 primary CRC samples (TCGA), 3 CRC cell lines (in house), and their matched normal samples. In total, we detected 1,129,208 somatic substitutions and 430,149 indels in the tumor enhancerome. The rates of mutation in the enhancerome were similar to those previously reported for the exome <u>(Lawrence et al. 2013)</u>, with a clear separation of hypermutator (10->100 mutations per Mb) and non-hypermutator samples (<1 – 10 mutations per Mb; Figure 3-figure supplement 1c). Of the 16 hypermutated samples, 10 had MMR-mutational signatures and were designated MSI, and 6 had POLE-mutational signatures (Alexandrov et al. 2013). Thirteen percent (46/355) of enhancer indels detected in the CRC cell lines were found to be recurrent in at least one of the 50 WGS samples. Fifteen indels were significantly recurrent in at least 2 CRC samples (Figure 3d, Figure 3-source data 2). The recurrent indels were found predominantly in MSI samples, and also included MSS samples with relatively high mutation rates, suggesting that enhancer activation by indels occurs whenever a sufficiently high mutation rate is reached. Furthermore, consistent with their positive selection, recurrent indels occurred more frequently in gained enhancers than in enhancers shared between tumor and normal crypts (Figure 3-figure supplement 1d). Genes associated with the most recurrent indels (4 of 50 samples) include *PIK3CB*, a catalytic subunit of phosphoinositide 3-kinase that has been previously implicated in breast cancer (Nakanishi et al. 2016), *USP8*, a deubiquitinase linked to EGFR signaling (Kim et al. 2017), and *MSX2*, a homeobox transcription factor associated with a diversity of growth-related functions (Satoh et al. 2008) (Figure 3d).

### MSI enhancer mutations recruit FOX transcription factors

We set out to identify transcription factors recruited to the indels and potentially responsible for enhancer activation. DNA motif scanning tools revealed forkhead (FOX) sites as most enriched at indels in gained enhancers (Table 1). FOX are known pioneer transcription factors in many cell types (Ang et al. 1993; Lupien et al. 2008; Iwafuchi-Doi et al. 2016), and are associated with enhancer re-programming (Pomerantz et al. 2015; Roe et al. 2017). FOX also scored as the top hit among factors predicted to bind with higher affinity to the indel than the wildtype sequence (Figure 4a-b, Figure 4-source data 1) and was the second most enriched motif (behind CREB) in the recurrent indels detected in the TCGA CRC cohort. Through integrative analyses of available FOX ChIP-seq datasets (Yan et al. 2013) with H3K27ac ChIP-seq data from LoVo cells, we determined that 83% (64/77) of gained enhancer indels show evidence of FOX enrichment by ChIP-seq. An example of an indel locus enriched for both FOXA2 and FOXO3 is shown in Figure 4c. Moreover, at the majority of FOX peaks with sufficient coverage at the indel to calculate allelic imbalance, FOX ChIP reads support allele-specific FOX binding (Figure 4d).

**Table 1:**
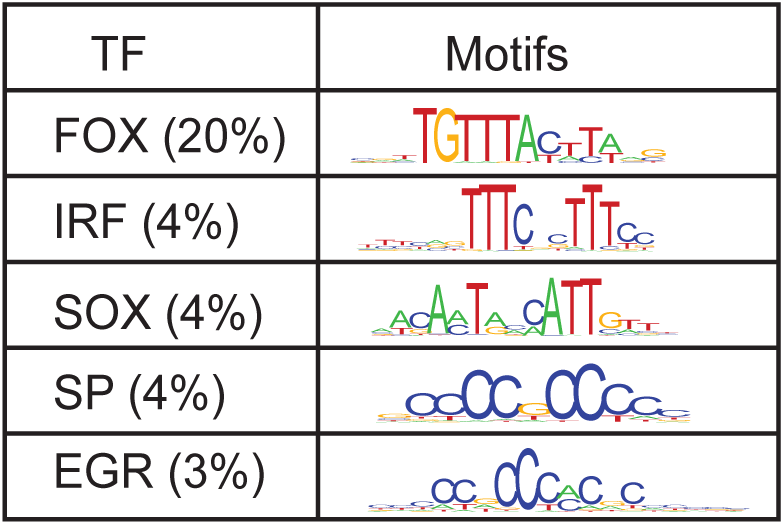
Top motifs enriched at indel loci. Motifs identified are from three computational programs, Deeptools, FIMO and Footprint. The logo displayed is from HOCOMOC v10 PWM’s for representative factors from each family (FOXC1, IRF1, SOX2, SP1, and EGR1). Total number of indels is 355.

**Figure 4:**
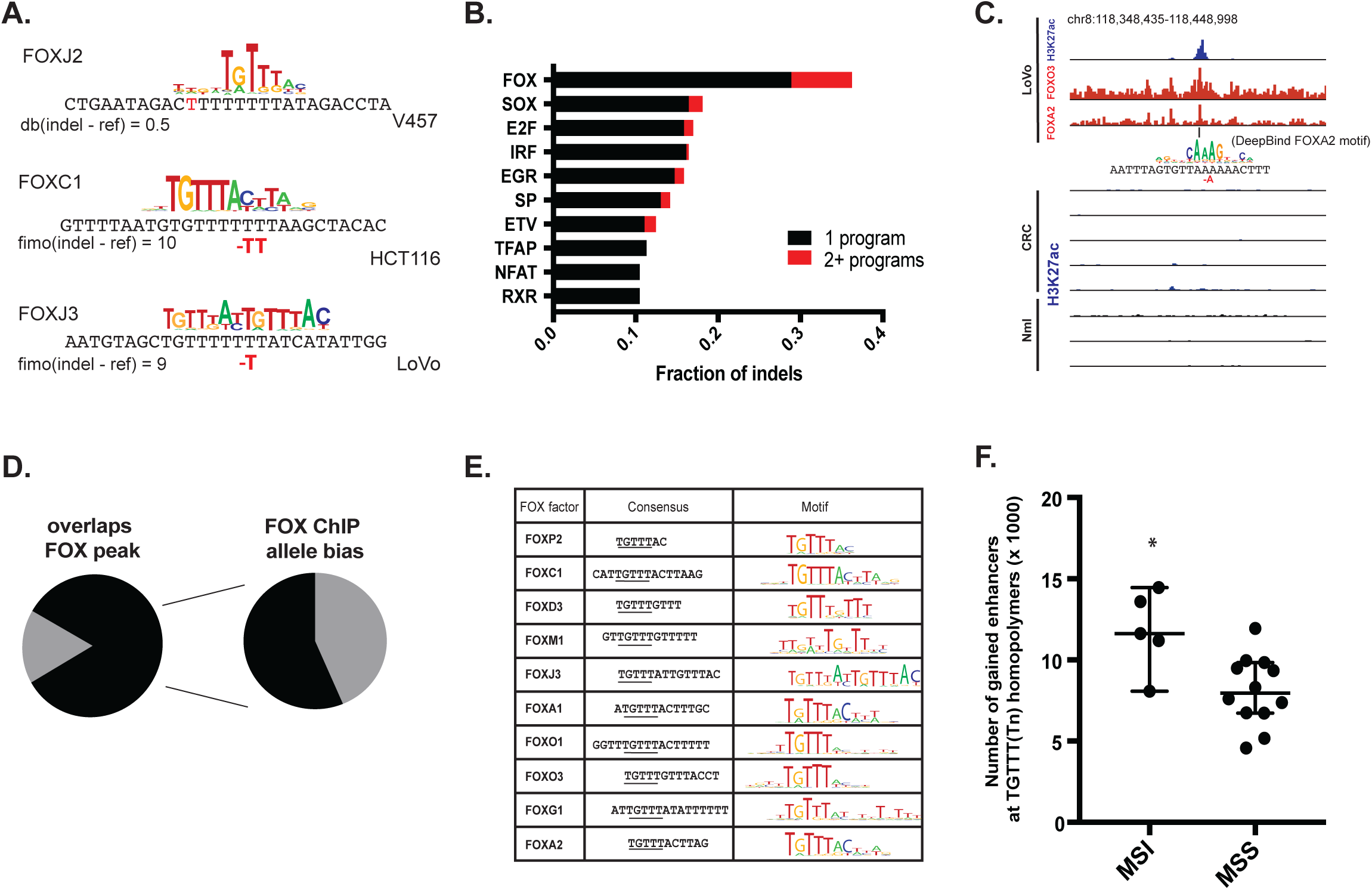
MSI enhancer indels recruit FOX factors. **A.** Examples of FOX motifs at MSI-indels, with change in affinity scores from computational programs. **B.** Fraction of indels predicted to increase transcription factor affinity by TF family, supported by one (black) or multiple (red) computational programs. (Figure 4-source data 1). **C.** Genome browser snapshot of MSI enhancer indel overlapping FOXA2 and FOXO3 peaks in LoVo cells (chr8:118,348,435-118,448,998). Motif of FOXA2 generated by DeepBind, which predicts increased binding of FOXA2 to the indel alelle, is shown. **D.** (left) Fraction of LoVo enhancer indels overlapping a FOX factor peak (64/77, or 83%). (right) Fraction of indels in FOX factor peaks and with >3 FOX ChIP read coverage showing allele-bias in FOX factor ChIP reads (11/19, or 58%). **E.** Motifs of FOX factors expressed in MSI lines and predicted to bind preferentially to indel alleles. **F.** Number of gained enhancers at TGTTT(Tn) motif in MSI versus MSS lines. * p <0.05, Wilcox ranksum test.

We examined the consensus motifs of FOX TFs expressed in MSI samples. The motifs are all A/T-rich sequences that closely resemble the ones most frequently mutated in MMR-deficient tumors. All share a core “TGTTT” within the consensus motif (Figure 4e, underlined). We hypothesized that if gained enhancers in MSI lines are formed by MMR-signature mutations that instigate FOX recruitment, then MSI samples should contain a higher frequency of gained enhancers at TGTTT(Tn) sequences than MSS samples. We retrieved all TGTTT(Tn) sequences across the genome and queried for gained H3K27ac enrichment. Strikingly, this unbiased analysis revealed that MSI samples had 50% more gained enhancers at TGTTT(Tn) sequences than MSS samples (Figure 4f). These observations indicate that gained enhancers arise more frequently at the sequences most prone to mutation as a result of MMR-deficiency. Coupled with the finding that these sites closely resemble the consensus motifs of FOX factors, the computational prediction that the indels increase FOX affinity, and the ChIP-seq results indicating allele-biased FOX binding to the indels, these observations suggest FOX factors play an important role in mediating enhancer activation at the indels. We note however, that given the degenerate nature of the FOX consensus motifs, our studies are limited in determining which specific FOX factor(s) is responsible for the activation.

### Induction of MSI phenotype yields enhancer mutations

To functionally test if mutations resulting from MMR deficiency can activate enhancers, we set out to introduce signature MMR mutations in CRC cells in a manner that recapitulates the natural way in which these mutations arise in MSI tumors. If they are indeed functional, introduction of the indels should lead to enhancer activation in combination with a bias in the H3K27ac ChIP-seq signal. We used CRISPR/Cas9 to knockout the *MLH1* gene in Colo-205, a microsatellite stable CRC cell line (workflow summarized in Figure 5a). Homozygous *MLH1* knockout was confirmed by sequencing and western blot (Figure 5b). Following 2.5 months of cell culture, subcloning, and expansion, several *MLH1*^*-*/-^ clones tested positive by PCR assay for the MSI phenotype (Figure 5c). Two *MLH1*^*-*/-^ and two parental wildtype clones were selected for H3K27ac ChIP-seq analysis. Applying our analysis pipeline, we uncovered enhancer indels matching the MMR-associated signature observed in the MSI lines of our panel, namely a high indel rate affecting homopolymers (10x the rate in the WT clones), a bias for short, 1-2 bp deletions, and a bias for poly-(A/T) tracts (Figure 5d, Figure 5-source data 1). Most indels in the *MLH1*^-/-^ cells (1357/1828, or 74%) arose in H3K27ac peaks that were already present at similar levels in parental wildtype clones. This was not unexpected, and likely reflects the switch from low to high mutation rate in open chromatin upon loss of MMR function, as observed previously in Figure 2a. Strikingly, we identified 45 indels in enhancers that showed at least a 1.5 fold increase in the H3K27ac signal in *MLH1*^-/-^ cells compared to parental wildtype cells (Figure 5e shows an example). We compared the percentage of mutant-allele reads at the 45 gained enhancer indels to that of the 1357 shared enhancer indels. Strikingly, the gained enhancer indels were more often allele-biased (Figure 5f) suggesting the basis of enhancer activation was acquisition of the indel. We further note that the frequency of indel-gained enhancer events detected in this experiment is likely to be lower than in naturally-derived MSI tumors, since the CRISPR-engineered cells spent a limited time in culture and were grown under conditions that do not recapitulate selective pressures of the tumor microenvironment.

**Figure 5:**
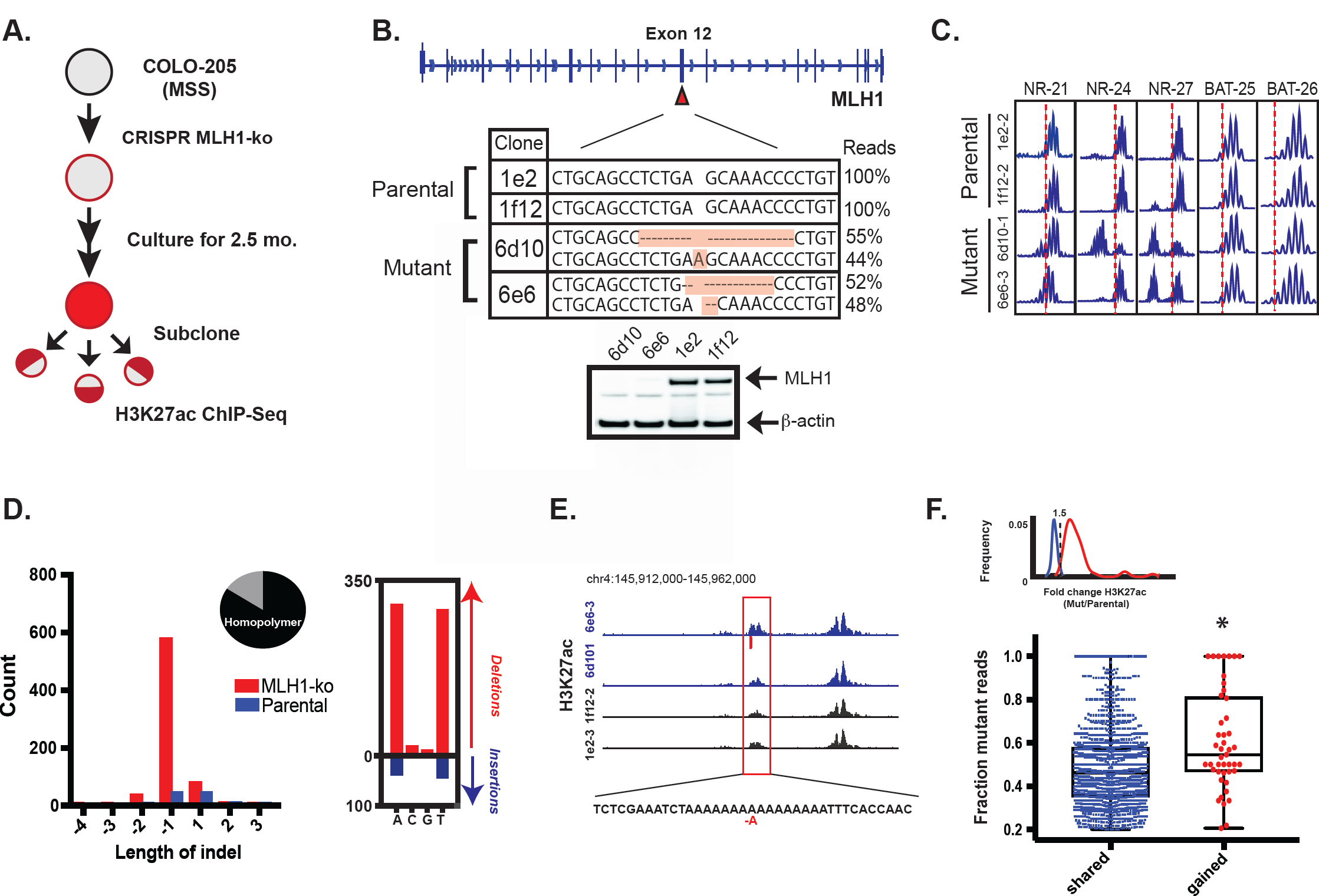
Induction of MSI phenotype yields enhancer mutations. **A.** Overview of MSI induction experiment. **B.** CRISPR-mediated knockout of *MLH1*. Nucleotides shaded in orange were determined by targeted NGS to be inserted/deleted. (bottom) Western blot analysis of MLH1 in *MLH1* wildtype and mutant clones. Beta-actin is shown as a loading control. **C.** PCR assay of five MSI markers (NR-21, NR-24, NR-27, BAT-25, and BAT-26) in *MLH1*^+/+^ and *MLH1*^-/-^clones after culturing for 2.5 months. Distribution of enhancer indel lengths. Pie chart represents fraction of indels affecting homopolymers (1317/1563, 84%). (right) Count of homopolymer insertion-(blue) and deletion-(red) mutations by mononucleotide repeat. **E.** Genome browser snapshot of indel (red bar) associated with increase in H3K27ac signal (chr4:145,912,000-145,962,000). Y-axis scales are all 0 to 90. **F.** Mutant allele fraction at indels in shared and gained H3K27ac peaks. (top) Density of H3K27ac signal fold change (*MLH1*^-/-^/*MLH1*^+/+^) for peaks defined as shared (<1.5 fold lesser or greater enrichment; blue) and gained (>1.5 fold greater enrichment; red) in *MLH1*^-/-^cells, relative to *MLH1*^+/+^ cells. (bottom) Dot plot of the mutant allele fraction distribution for indels in shared (n = 1357) and gained (n = 45) peaks. * p < 0.001, Wilcox ranksum test. (Figure 5-source data 1).

### MSI enhancer indels are propagated in tumors

To test if loss of *MLH1* induces indels that activate enhancers *in vivo*, we introduced *MLH1* knockout (clone 6e6-3) and wildtype (clone 1f12-2) cells into mice via intrasplenic injection (workflow in summarized in Figure 6a). In this assay, tumor cells typically form clonal liver metastases, but are also known to form peritoneal tumors (Lee et al. 2014). To simulate different micro-environmental pressures, we used two mouse strains (nude and NSG) with different levels of immune competence. Three months post-injection, 5 liver tumors were harvested from mice seeded with *MLH1*^*-*/-^ cells (4 tumors from one nude mouse and 1 tumor from one NSG mouse). Two peritoneal tumors were harvested from one NSG mouse injected with *MLH1*^+/+^ cells. We then performed H3K27ac ChIP-seq profiling of all 7 tumors. Unsupervised cluster analysis of the enhancer landscapes separated *MLH1*^+/+^ from *MLH1*^-/-^ tumors, and there was remarkable consistency among tumors with matched genotypes (Figure 6b). Analysis of the mutations detected from H3K27ac ChIP-seq of the *MLH1*^-/-^ tumors again revealed predominantly small 1-2 bp deletions in mononucleotide tracts of A/T repeats (Figure 6c, Figure 6-source data 1). We also noted a shift in the proportion of non-homopolymer mutations, including indels in larger tandem repeats and in non-tandem repeat regions. In each *MLH1*^-/-^ tumor, 6-10% of the indels were located in enhancers that showed at least a 1.5 fold increase in the H3K27ac signal relative to wildtype tumor cells. Consistent with an enhancer-activating role, indels in gained peaks again showed higher mutant allele fractions compared to indels in H3K27ac peaks already present at similar levels in the parental wildtype cells (Figure 6c, bottom). Using the ChIP-seq data from all 7 tumors, we identified 20 indels in *MLH1*^*-*/-^ tumors that showed a perfect correlation with an increase in H3K27ac, like the two examples shown in Figure 6d. Strikingly, these 20 indels also showed high mutant allele fractions (Figure 6e). Together, these data provide further functional support that MSI signature mutations activate enhancers *in vivo*.

**Figure 6:**
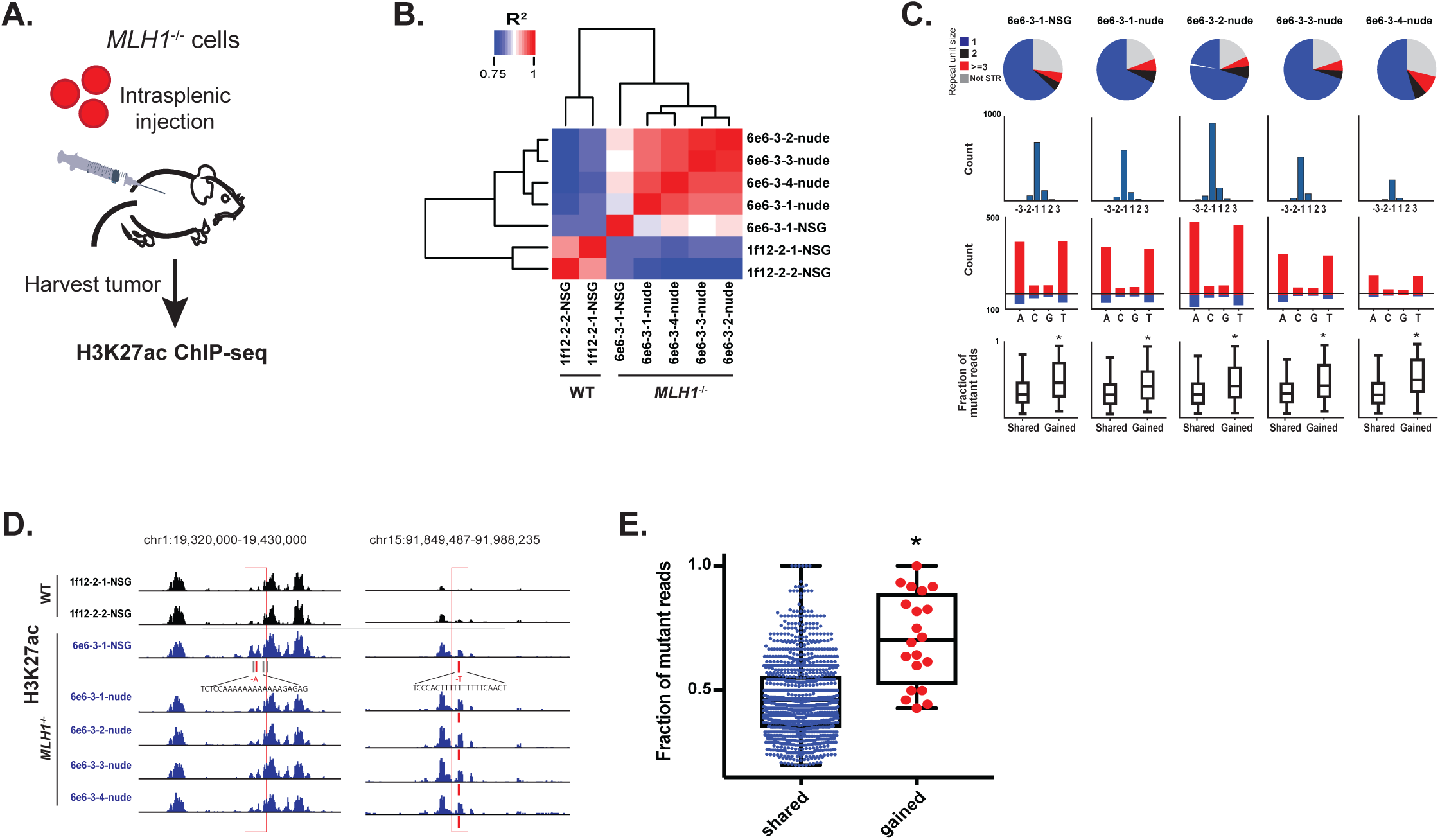
MSI enhancer indels are propagated in tumors. **A.** Overview of mouse tumor formation assay. Heatmap of R^2^ values from Pearson correlation of H3K27ac signal at enhancers, for pairs of tumors. (top) Pie charts show distribution of types of mutations (mononucleotide, dinucleotide, trinucleotide or higher, and non-short tandem repeat). (2nd row) Histogram of indel sizes. (3rd row) Bar plots of homopolymer indel frequency by mononucleotide repeat. (bottom) Boxplots of the mutant allele fraction distribution for indels in gained peaks and peaks shared with wildtype tumors. * p < 0.001, Wilcox ranksum test. Genome browser snapshots of indels (red bars) detected in *MLH1*^-/-^tumors and associated with increased H3K27ac signal (left: chr1:19,320,000-19,430,000. Y-axis scales are all 0-156; right: chr15:91,849,487-91,988,235. Y-axis scales are all 0-95). **E.** Dot plot of the mutant allele fraction distribution for indels in shared (n= 1288) peaks and indels correlating with gained H3K27ac enrichment (n= 20). * p < 0.001, Wilcox ranksum test. See Figure 6-source data 1.

## Discussion

The search for functional mutations in non-coding regions of tumor genomes is an area of intense investigation. Beyond those affecting *TERT, ER, FOXA1* and others, regulatory mutations are presumed rare and many are considered passengers (Fredriksson et al. 2014; Cuykendall et al. 2017). Consistent with this supposition, the majority of samples in our CRC cohort showed a low enhancer mutation rate. However, a clear exception are CRC tumors whose mutation rates across the genome are extraordinarily high, like the MSI subtype. In MSI CRC samples, we detected a large number of indel mutations that correlate with higher levels of H3K27ac in tumor cells relative to normal colon cells. In addition to showing allele-bias, several lines of evidence indicate these indels functionally activate enhancers and are not merely random passengers. First, they show signatures of positive selection. Second, their target genes, in addition to being enriched for cancer-related functions, are more highly expressed when both the gained enhancer and indel are present compared to when only the gained enhancer is present. Third, a higher-than-expected number of the enhancer indels are recurrent across primary CRC tumor samples. Fourth, the indels are predicted to enhance FOX binding and an unbiased genome-wide scan of the core FOX motif sequence TGTTT(Tn) showed 50% more gained H3K27ac signals at this motif in MSI samples than MSS samples. Lastly, introduction of the indels via CRISPR-Cas9 inactivation of *MLH1*-thereby mimicking the natural way in which these mutations occur - led to allele-biased enhancer activation and selection of enhancers both in cell culture and in full blown tumors, suggesting that the basis of enhancer activation in these cells was acquisition of the mutation. Based on these findings, we conclude that in CRC, deficiencies in mismatch repair lead to the appearance of indels that are recognized by FOX factors, which turn these sites into functional cis-regulatory elements. It has long been established that signature MMR-mutations, when they arise in coding regions, can impact gene splicing or shift the reading frame. Indel mutations that induce a frameshift in *TGFBR2* are a notable example (Markowitz et al. 1995). Our studies here expand the repertoire of functional MSI-type mutations to non-coding regions, and offer a new method of identifying functional mutations in regulatory elements.

There are notable differences between the functional regulatory mutations described here and those previously linked to *TERT, ER, TAL1, MYC*, and *FOXA1*. The latter are considered tumor-initiating events and activate *bona fide* drivers of oncogenesis. In contrast, most of the enhancer mutations reported here are a consequence of MMR-inactivation and likely arose subsequent to mutations in canonical CRC drivers like *APC, TP53*, and *KRAS*. While some of the enhancer indel targets could be oncogenes, most are genes with cancer-related functions that likely provide a selective growth advantage. Another key difference is the high occurrence of enhancer mutations in MSI CRC versus other cancers. Given their prevalence and widespread effects on gene expression, MSI enhancer mutations could be considered “reprogrammers” of cell identity. We further note, given that MSI is a continuously evolving phenotype, that enhancer mutations due to MMR-deficiencies could play an important role in tumor evolution and the emergence of drug resistance. Furthermore, as the MSI subset of CRC is a particularly good candidate for immunotherapy, it would be worth investigating if enhancer mutations instigate expression of PD-1 pathway genes or neoantigen-producing genes that contribute to tumor immunogenicity and may therefore be exploited for refining response prediction.

## Methods

### ChIP-seq

H3K27ac and H3K4me1 ChIP-seq datasets from CRC cell lines and primary tumors are previously described in our previous publications (Cohen et al. 2017; Akhtar-Zaidi et al. 2012) and available in GEO (Accession numbers GSE36401 and GSE77737). H3K27ac ChIP-seq was performed on LoVo, Colo-205, and MLH1^-/-^ Colo-205 cells as previously described. ChIP data processing, alignment, peak-calling, and identification of differentially enriched-peaks relative to normal colonic crypts were done as previously described.

### Detection of enhancer mutations

H3K27ac ChIP seq reads were aligned to the human genome (hg19) with Bowtie2. Reads were realigned around regions with evidence of indels (>1 indel supporting read) using GATK v.2-2-gec30cee. A custom FASTA file was created incorporating the candidate indel sequences and their flanking regions such that the length of the flanking region equals the length of the longest aligned read. Any read that aligned perfectly to both reference and custom indel genomes was discarded. Samtools 1.2 was used to generate a multi-way pileup output file for each filtered BAM. Indels were called using VarScan.v2.4.0 from the pileup output, requiring at least 10X coverage and 20% of reads supporting the indel. To exclude possible germline variants, indels matching dbSNPs from the 1000 Genomes Project and/or SNP142 indels were filtered out.

Candidate enhancer-activating mutations were prioritized based on whether they correlate with gained H3K27ac enrichment. Binary matrices of indels and RPKM matrices were constructed for all H3K27ac peaks. A peak was reported as correlating with an indel if the following conditions were met:

1) Peak RPKM for the sample the peak was called in is at least 2.

2) Minimum peak RPKM for samples with indels in the peak is greater than the maximum RPKM for the samples with no reported indels in the same peak.

3) No reads support the indel in samples not called to have the indel

Indels were filtered if they overlapped artifact regions in the consensus Blacklist (https://sites.google.com/site/anshulkundaje/projects/blacklists). An empirical approach was used to identify indel calls that are likely alignment errors. For each indel, the sequence 50 bp upstream and downstream was aligned to the human genome (Blat, from UCSC Genome Browser). The reference allele was replaced with the indel allele, to simulate the alignment of indel-supporting ChIP reads. Indels whose second-highest alignment score was > 50 indicated potential alignment error and were discarded.

Indels with imbalanced read distributions favoring the indel allele were prioritized, as this suggests a scenario whereby the enhancer signal and indel co-occur on the same allele. Bias in the number of reads supporting the mutation was quantified by the complement of the cumulative binomial distribution (upper-tail), with a probability of success of 0.5. Multiple test correction was performed using the Benjamini-Hochberg method controlling the FDR at 0.2.

### Correlation of H3K27ac enrichment and indel rate

Chromosome 14 was split into 537 bins of 200 kb. For each bin, the median H3K27ac signal from the V410 or HCT-116 bigWig was retrieved. Indel calls from either MSS samples or MSI samples in the TCGA cohort were pooled and indels were counted in each bin. The H3K27ac signal and indel rate were quantile-normalized and smoothed. For comparisons of the indel rate in gained and shared enhancers, the indel rate was defined as the number of indels divided by the length of the enhancer peak, and plotted as a function of the peak’s median H3K27ac bigWig signal. Regression analysis using a generalized linear model (binomial distirbution) was performed to get the significance of gained enhancer status as a predictor for indel rate.

### Prediction of indel gained enhancer gene targets

GREAT (version 3.3.0) was used to pair gene targets with enhancers at each locus (“basal plus extension” setting), and to get the distance to the nearest transcription start site. For each predicted target gene, the (quantile-normalized) expression from microarray data was retrieved for the line harboring the indel gained enhancer, and lines harboring gained enhancers without indels at the locus. Fold change relative to the median expression of that gene in 5 normal crypt lines was calculated. Enriched gene ontology terms were obtained by inputting GREAT genes (with > 2 fold increase in expression relative to normal crypt) to the GSEA web tool to compute overlap with MSigDB database of gene-sets. Towards a comprehensive list of genes with function in cancer, gene lists from various sources, including the cancer gene census (COSMIC), amplified genes in cancer (Santarius et al. 2010), and genes in CRC pathways (KEGG cancer pathways, KEGG CRC, KEGG MAPK, and KEGG WNT) or involved in cell proliferation (GO positive regulation of cell proliferation), were compiled.

### Mutation calling from TCGA COAD and in-house WGS data

Whole genome sequence reads aligned to GRCh37 (.bam files) of tumor and normal matched pairs were downloaded from the Cancer Genomics Hub (UCSC) using the GeneTorrent tool, and re-processed according to GATK best practices. First, base quality scores were reverted (Picard v.1.104 RevertSam), but duplicate and alignment information was retained. Next, the alignment of reads around indels was redone and then base quality scores were re-calibrated using GATK IndelRealinger and BaseRecalibrator (version 3.2-2-gec30cee), respectively. Somatic substitutions and indels were called using 2 mutation callers, MuTect2 (v3.5-0-g3628e4) and Varscan2 (v2.3.9) using default settings, and the intersection of the calls was used for further analysis. Variants in common dbSNP and in 1000 Genomes were filtered out as likely germline events. Significance of the recurrence distribution was determined by randomly picking indels (same number as actual recurrent indel set) from the binary matrix and finding the average recurrence distribution of those random indels.

### Transcription factor motif analysis

Ten (non-homopolymer) base pairs of the reference genome (hg19) upstream and downstream of the indel were retrieved and used to flank either the reference allele or the indel allele. Both indel and reference sequences were input to 3 computational programs which score the affinity of their interaction with transcription factors: Deepbind (Alipanahi et al. 2015), FIMO (Grant et al. 2011) with the HOCOMOCO v.10 motif database, and Footprint (Sebastian and Contreras-Moreira 2014), with the Human-TF v2.0 (Jolma et al. 2015) motif database. For each program, a “binding score” was retrieved for each TF-sequence pair, and compared between reference and indel alleles. A prediction of increased binding was counted if binding score_indel_ > binding score_reference_ and the predicted TF is expressed in the cell line harboring the indel. TF predictions were collapsed by TF family. Unless otherwise noted, logos were created from HOCOMOCO v.10 PWM’s. FOX ChIP-seq data from the LoVo cell line (Yan et al. 2013) was downloaded (GEO GSE51142). Reads were re-aligned to the human reference genome (hg19) using Bowtie2 to generate BAM files. Peaks were called using macs1.4. Reads at indel loci were retrieved using samtools view command. To calculate allele bias, indels with coverage of 4 or more FOX ChIP-seq reads were selected, and the complement of the cumulative binomial distribution, with a probability of success of 0.5, was used. The false discovery rate was set to 0.2.

### CRISPR-mediated MLH1-knockout in COLO-205 cells

*MLH1* knockout was performed in collaboration with the Genome Engineering and iPSC Center (GEiC) at Washington University. Guide RNAs were designed to target a conserved exon (exon 12) of the *MLH1* gene. The gRNAs and Cas9 were nucleofected into Colo-205 (MSS) cells, and then individual lines were subcloned. CRISPR-induced indels were verified by targeted NGS. WT controls were unmodified clones from the same nucleofected pool of cells.

### Cell culture

*MLH1* knockout and parental WT lines were cultured for 2 months in RPMI-1640 media supplemented with 10% fetal bovine serum (ThermoFisher 11875093) to accumulate mutations. Monoclonal cells were then derived by limiting dilution, assayed for instability at MSI markers, and then expanded for 3-4 weeks before using in H3K27ac ChIP-seq experiment.

### PCR assay for microsatellite instability

Genomic DNA was extracted and PCR-amplified at five MSI markers (Buhard et al. 2004; Buhard et al. 2006). PCR primers are from Buhard et al. 2006, with one primer in each pair labeled with a fluorescent dye for combined analysis. PCR products were run on a Genetic Analyzer ABI 3730 to produce fragment profiles, which were visualized using Peak Scanner version 1.0 (ThermoFisher).

### Mice

Two 6-week old female Nod-Scid IL-2Rg-/-(NSG) mice and two nude mice were each injected intra-splenically with 10^6^ (6e6-3 cells or 1f12-2) cells. After 3 months, mice were sacrificed. Tumors were harvested and homogenized for H3K27ac ChIP-seq experiment. All mouse experiments were done according to Institutional Animal Care and Use Committees (IACUC) guidelines.

## Acknowledgements

We thank Grace Lee from Dr. David Wald’s group for experimental assistance preparing the mice for surgery. We thank the Case Western Reserve University Genomics Core for their support with ChIP-sequencing and genotyping, and the High Performance Computing Cluster for their computational infrastructure. We thank Samuel Li and Sneha Grandhi from Dr. Thomas LaFramboise’s lab for help downloading WGS data. We thank Shiyi Yin from Dr. Ann Harris’ lab and Chen Weng from Dr. Fulai Jin’s lab for valuable discussions. Finally, we thank Monica Sentmanat and the Genome Engineering and iPSC Center at Washington University at St. Louis for generating the MLH1^-/-^ clones.

---

**Figure 1-figure supplement 1.**
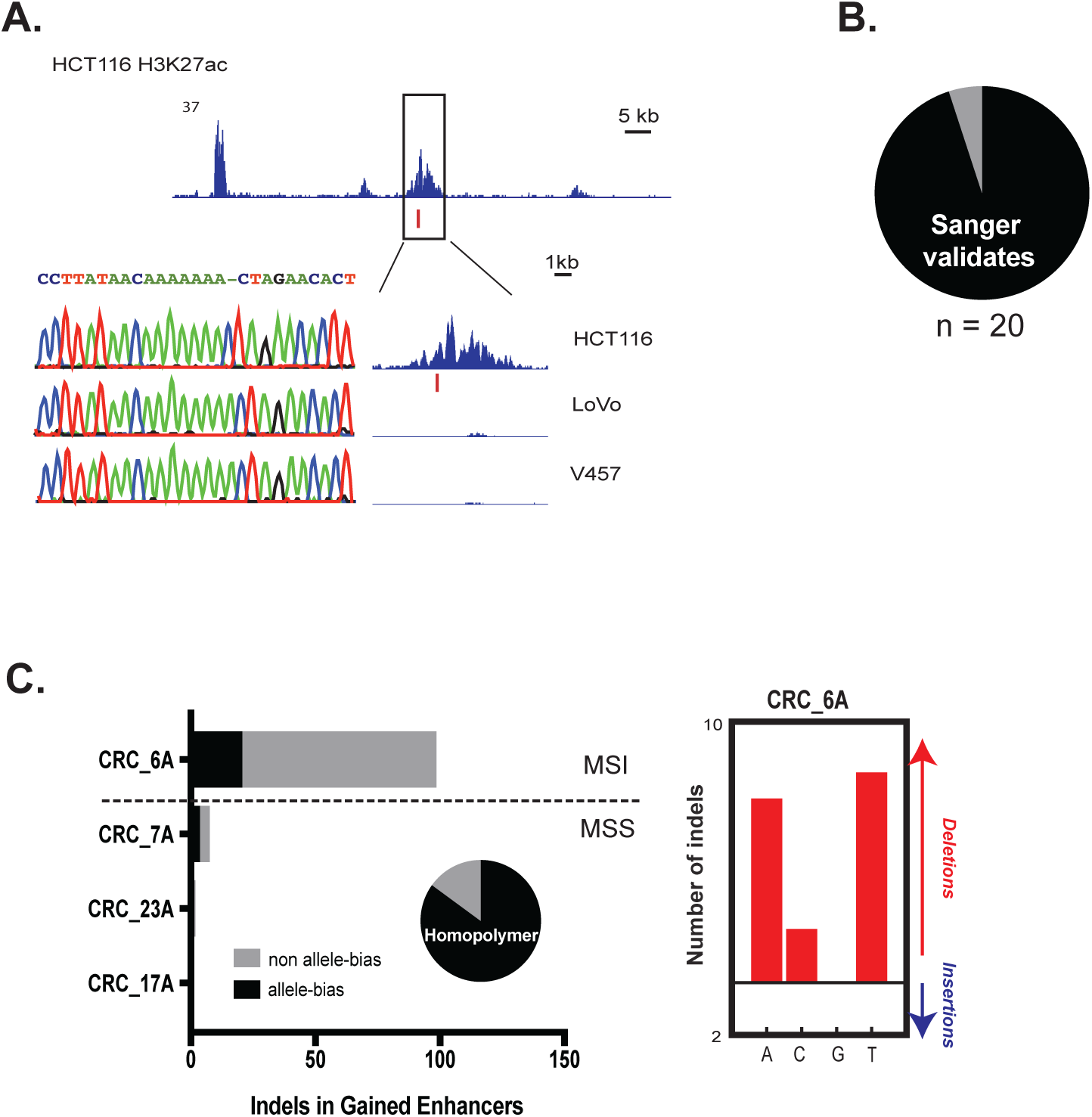
Validation of enhancer indels. **A.** Example of a gained enhancer indel confirmed by Sanger sequencing. (top) Genome Browser snapshot of a putative enhancer-activating indel detected in HCT-116 cells (chr5:1,870,785-1,999,125). (bottom) Sanger traces of the locus and zoomed in view of the peak (chr5:1,928,000-1,942,000) for HCT-116 cells, and 2 non-indel cell lines (LoVo and V457). Y-axis scales are all 0 to 24. **B.** Fraction of enhancer indels (19/20) validated by Sanger sequencing. (left) Number of enhancer indels detected in four primary tumors. Pie chart shows fraction of indels detected in the CRC_6A sample that lie in homopolymers (17/20, or 85%). (right) Number of indels in homopolymers of A,C,G, or T in the CRC_6A sample.

**Figure 3-figure supplement 1.**
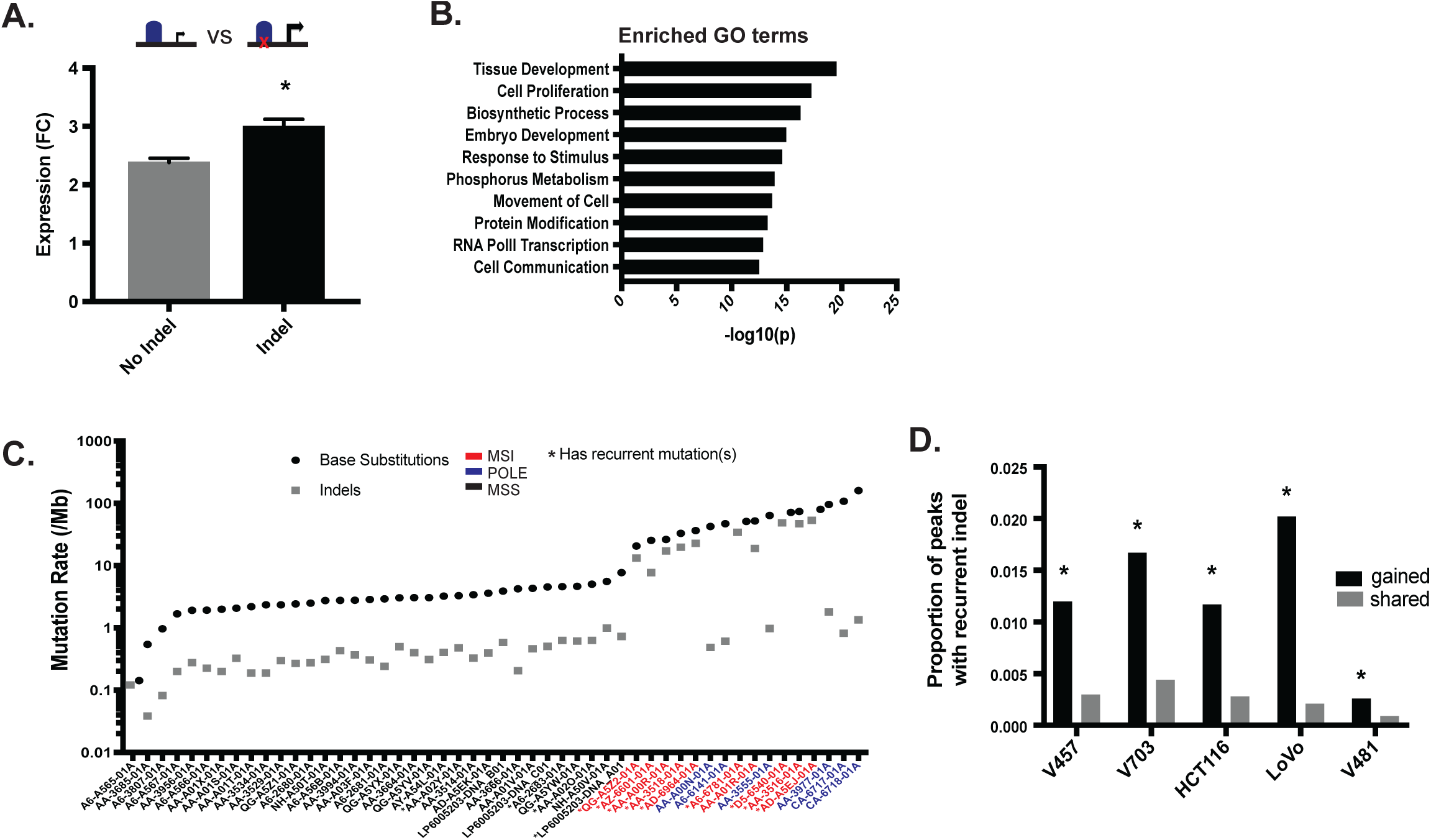
**A.** Mean fold-change of expression (CRC/crypts) of genes corresponding to all gained enhancers containing indels (n = 10,971 gene-sample pairs; grey), versus the same genes in cell lines with gained enhancers but lacking indels (n= 58,412 gene-sample pairs; black). * p< 0.05, 2-sample t-test. **B.** Enriched molecular function (GO) terms for genes associated with indel gained enhancers, ranked by significance. **C.** Rate of substitutions and indels detected in cohort of 50 primary CRC samples, ordered by total mutation rate. **D.** Proportion of gained enhancers (black) and enhancers shared with normal crypt (grey) with at least 1 indel recurrent in primary CRC tumors. * p<0.05, Z-test of proportions.

## References

Akhtar-Zaidi, Batool, Richard Cowper-Sal-lari, Olivia Corradin, Alina Saiakhova, Cynthia F. Bartels, Dheepa Balasubramanian, Lois Myeroff, et al. 2012. “Epigenomic Enhancer Profiling Defines a Signature of Colon Cancer; PMC3711120.” Science 336 (6082): 736–739. doi:10.1126/science.1217277. http://dx.doi.org/10.1126/science.1217277whttps://www.ncbi.nlm.nih.gov/pubmed/22499810 https://www.ncbi.nlm.nih.gov/pmc/articles/PMC3711120whttp://www.sciencemag.org/cgi/pmidlookup?view=long&pmid=22499810.

Alexandrov, Ludmil B., Serena Nik-Zainal, David C. Wedge, A. J. Samuel, Sam Behjati, Andrew V. Biankin, Graham R. Bignell, et al. 2013. “Signatures of Mutational Processes in Human Cancer.” Nature 500 (7463): 415–421. doi:10.1038/nature12477. http://dx.doi.org/10.1038/nature12477http://www.nature.com/nature/journal/v500/n7463/full/nature12477.html.

Alipanahi, Babak, Andrew Delong, Matthew T. Weirauch, and Brendan J. Frey. 2015. “Predicting the Sequence Specificities of DNA-and RNA-Binding Proteins by Deep Learning.” Nat.Biotechnol. 33: 831. doi:10.1038/nbt.3300. http://dx.doi.org/10.1038/nbt.3300whttps://www.nature.com/articles/nbt.3300.

Ang, S. L., A. Wierda, D. Wong, K. A. Stevens, S. Cascio, J. Rossant, and K. S. Zaret. 1993. “The Formation and Maintenance of the Definitive Endoderm Lineage in the Mouse: Involvement of HNF3/Forkhead Proteins.” Development 119 (4): 1301–1315. http://dev.biologists.org/content/119/4/1301whttps://www.ncbi.nlm.nih.gov/pubmed/8306889.

Baross-Francis, Agnes, Naila Makhani, R. M. Liskay, and Frank R. Jirik. 2001. “Elevated Mutant Frequencies and Increased C : G→T : A Transitions in Mlh1-/-Versus Pms2-/-Murine Small Intestinal Epithelial Cells.” Oncogene 20: 619. doi:10.1038/sj.onc.1204138. http://dx.doi.org/10.1038/sj.onc.1204138whttps://www.nature.com/articles/1204138.

Battey, J., C. Moulding, R. Taub, W. Murphy, T. Stewart, H. Potter, G. Lenoir, and P. Leder. 1983. “The Human C-Myc Oncogene: Structural Consequences of Translocation into the IgH Locus in Burkitt Lymphoma.” Cell 34 (3): 779–787. https://www.ncbi.nlm.nih.gov/pubmed/6414718https://linkinghub.elsevier.com/retrieve/pii/0092-8674(83)90534-2.

Beroukhim, Rameen, Xiaoyang Zhang, and Matthew Meyerson. 2016. “Copy Number Alterations Unmasked as Enhancer Hijackers.” Nat.Genet. 49 (1): 5–6. doi:10.1038/ng.3754. http://dx.doi.org/10.1038/ng.3754https://www.ncbi.nlm.nih.gov/pubmed/28029156.

Buhard, Olivier, Francesca Cattaneo, Yick Fu Wong, So Fan Yim, Eitan Friedman, Jean-François Flejou, Alex Duval, and Richard Hamelin. 2006. “Multipopulation Analysis of Polymorphisms in Five Mononucleotide Repeats used to Determine the Microsatellite Instability Status of Human Tumors.” J.Clin.Oncol.24 (2): 241–251. doi:10.1200/JCO.2005.02.7227. http://dx.doi.org/10.1200/JCO.2005.02.7227https://www.ncbi.nlm.nih.gov/pubmed/16330668 http://ascopubs.org/doi/abs/10.1200/JCO.2005.02.7227?url_ver=Z39.88-2003&rfr_id=ori:rid:crossref.org&rfr_dat=cr_pub%3dpubmed.

Buhard, Olivier, Nirosha Suraweera, Aude Lectard, Alex Duval, and Richard Hamelin. 2004. “Quasimonomorphic Mononucleotide Repeats for High-Level Microsatellite Instability Analysis; PMC3888729.” Dis.Markers 20 (4-5): 251–257. https://www.ncbi.nlm.nih.gov/pubmed/15528790https://www.ncbi.nlm.nih.gov/pmc/articles/PMC3888729.

Cohen, Andrea J., Alina Saiakhova, Olivia Corradin, Jennifer M. Luppino, Katreya Lovrenert, Cynthia F. Bartels, James J. Morrow, et al. 2017. “Hotspots of Aberrant Enhancer Activity Punctuate the Colorectal Cancer Epigenome.” Nat.Commun. 8: 14400. doi:10.1038/ncomms14400. http://dx.doi.org/10.1038/ncomms14400 http://www.nature.com/articles/ncomms14400.

Cuykendall, Tawny N., Mark A. Rubin, and Ekta Khurana. 2017. “Non-Coding Genetic Variation in Cancer.” Current Opinion in Systems Biology 1: 9–15. doi:10.1016/j.coisb.2016.12.017. http://www.sciencedirect.com/science/article/pii/S2452310017300094http://dx.doi.org/10.1016/j.coisb.2016.12.017 https://www.sciencedirect.com/science/article/pii/S2452310017300094.

Duerr, Eva-Maria, Yusuke Mizukami, Kentaro Moriichi, Manish Gala, Won-Seok Jo, Hirotoshi Kikuchi, Ramnik J. Xavier, and Daniel C. Chung. 2012. “Oncogenic KRAS Regulates BMP4 Expression in Colon Cancer Cell Lines; PMC3362092.” Am.J.Physiol.Gastrointest.Liver Physiol. 302 (10): 1223. doi:10.1152/ajpgi.00047.2011. http://dx.doi.org/10.1152/ajpgi.00047.2011 https://www.ncbi.nlm.nih.gov/pubmed/22383492 https://www.ncbi.nlm.nih.gov/pmc/articles/PMC3362092 http://www.physiology.org/doi/abs/10.1152/ajpgi.00047.2011?url_ver=Z39.88-2003&rfr_id=ori:rid:crossref.org&rfr_dat=cr_pub%3dpubmed https://www.physiology.org/doi/10.1152/ajpgi.00047.2011.

Fredriksson, Nils J., Lars Ny, Jonas A. Nilsson, and Erik Larsson. 2014. “Systematic Analysis of Noncoding Somatic Mutations and Gene Expression Alterations Across 14 Tumor Types.” Nat.Genet. 46 (12): 1258–1263. doi:10.1038/ng.3141. http://dx.doi.org/10.1038/ng.3141https://www.ncbi.nlm.nih.gov/pubmed/25383969.

Grant, Charles E., Timothy L. Bailey, and William Stafford Noble. 2011. “FIMO: Scanning for Occurrences of a Given Motif; PMC3065696.” Bioinformatics 27 (7): 1017–1018. doi:10.1093/bioinformatics/btr064. http://dx.doi.org/10.1093/bioinformatics/btr064 https://www.ncbi.nlm.nih.gov/pubmed/21330290 https://www.ncbi.nlm.nih.gov/pmc/articles/PMC3065696 https://academic.oup.com/bioinformatics/article-lookup/doi/10.1093/bioinformatics/btr064 https://www.ncbi.nlm.nih.gov/pmc/articles/PMC3065696/.

Hegan, Denise Campisi, Latha Narayanan, Frank R. Jirik, Winfried Edelmann, R. M. Liskay, and Peter M. Glazer. 2006. “Differing Patterns of Genetic Instability in Mice Deficient in the Mismatch Repair Genes Pms2, Mlh1, Msh2, Msh3 and Msh6; PMC2612936.” Carcinogenesis 27 (12): 2402–2408. doi:10.1093/carcin/bgl079. http://dx.doi.org/10.1093/carcin/bgl079 https://www.ncbi.nlm.nih.gov/pubmed/16728433 https://www.ncbi.nlm.nih.gov/pmc/articles/PMC2612936 https://academic.oup.com/carcin/article-lookup/doi/10.1093/carcin/bgl079.

Hnisz, Denes, Abraham S. Weintraub, Daniel S. Day, Anne-Laure Valton, Rasmus O. Bak, Charles H. Li, Johanna Goldmann, et al. 2016. “Activation of Proto-Oncogenes by Disruption of Chromosome Neighborhoods.” Science: aad9024. doi:10.1126/science.aad9024. http://science.sciencemag.org/content/early/2016/03/02/science.aad9024 http://dx.doi.org/10.1126/science.aad9024 https://www.ncbi.nlm.nih.gov/pubmed/26940867.

Huang, Franklin W., Eran Hodis, Mary Jue Xu, Gregory V. Kryukov, Lynda Chin, and Levi A. Garraway. 2013. “Highly Recurrent TERT Promoter Mutations in Human Melanoma; PMC4423787.” Science 339 (6122): 957–959. doi:10.1126/science.1229259. http://dx.doi.org/10.1126/science.1229259https://www.ncbi.nlm.nih.gov/pubmed/23348506 https://www.ncbi.nlm.nih.gov/pmc/articles/PMC4423787 http://www.sciencemag.org/cgi/pmidlookup?view=long&pmid=23348506 http://science.sciencemag.org/content/339/6122/957.

Ionov, Y., M. A. Peinado, S. Malkhosyan, D. Shibata, and M. Perucho. 1993. “Ubiquitous Somatic Mutations in Simple Repeated Sequences Reveal a New Mechanism for Colonic Carcinogenesis.” Nature 363 (6429): 558–561. doi:10.1038/363558a0. http://dx.doi.org/10.1038/363558a0https://www.ncbi.nlm.nih.gov/pubmed/8505985.

Iwafuchi-Doi, Makiko, Greg Donahue, Akshay Kakumanu, Jason A. Watts, Shaun Mahony, B. Franklin Pugh, Dolim Lee, Klaus H. Kaestner, and Kenneth S. Zaret. 2016. “The Pioneer Transcription Factor FoxA Maintains an Accessible Nucleosome Configuration at Enhancers for Tissue-Specific Gene Activation.” Mol.Cell 62 (1): 79–91. doi:10.1016/j.molcel.2016.03.001. https://www.cell.com/molecular-cell/abstract/S1097-2765(16)00179-9 http://dx.doi.org/10.1016/j.molcel.2016.03.001 https://www.ncbi.nlm.nih.gov/pubmed/27058788.

Jolma, Arttu, Yimeng Yin, Kazuhiro R. Nitta, Kashyap Dave, Alexander Popov, Minna Taipale, Martin Enge, Teemu Kivioja, Ekaterina Morgunova, and Jussi Taipale. 2015. “DNA-Dependent Formation of Transcription Factor Pairs Alters their Binding Specificity.” Nature 527: 384. doi:10.1038/nature15518. http://dx.doi.org/10.1038/nature15518 https://www.nature.com/articles/nature15518.

Kandoth, Cyriac, Michael D. McLellan, Fabio Vandin, Kai Ye, Beifang Niu, Charles Lu, Mingchao Xie, et al. 2013. “Mutational Landscape and Significance Across 12 Major Cancer Types.” Nature 502 (7471): 333–339. doi:10.1038/nature12634. http://dx.doi.org/10.1038/nature12634http://www.nature.com/articles/nature12634.

Kim, Tae-Min, Peter W. Laird, and Peter J. Park. 2013. “The Landscape of Microsatellite Instability in Colorectal and Endometrial Cancer Genomes.” Cell 155 (4): 858–868. doi:10.1016/j.cell.2013.10.015. http://www.cell.com/abstract/S0092-8674(13)01291-9 http://dx.doi.org/10.1016/j.cell.2013.10.015 https://www.ncbi.nlm.nih.gov/pubmed/24209623.

Kim, Yunjung, Aya Shiba-Ishii, Tomoki Nakagawa, Ryan Edbert Husni, Shingo Sakashita, Tomoyo Takeuchi, and Masayuki Noguchi. 2017. “Ubiquitin-Specific Protease 8 is a Novel Prognostic Marker in Early-Stage Lung Adenocarcinoma.” Pathol.Int. 67 (6): 292–301. doi:10.1111/pin.12546. http://dx.doi.org/10.1111/pin.12546 https://www.ncbi.nlm.nih.gov/pubmed/28544031.

Latres, Esther, Roberto Chiarle, Brenda A. Schulman, Nikola P. Pavletich, Angel Pellicer, Giorgio Inghirami, and Michele Pagano. 2001. “Role of the F-Box Protein Skp2 in Lymphomagenesis.” Proc.Natl.Acad.Sci.U.S.A. 98 (5): 2515–2520. doi:10.1073/pnas.041475098. http://www.pnas.org/content/98/5/2515http://dx.doi.org/10.1073/pnas.041475098 https://www.ncbi.nlm.nih.gov/pubmed/11226270.

Lawrence, Michael S., Petar Stojanov, Craig H. Mermel, James T. Robinson, Levi A. Garraway, Todd R. Golub, Matthew Meyerson, Stacey B. Gabriel, Eric S. Lander, and Gad Getz. 2014. “Discovery and Saturation Analysis of Cancer Genes Across 21 Tumour Types.” Nature 505: 495. doi:10.1038/nature12912. http://dx.doi.org/10.1038/nature12912 https://www.nature.com/articles/nature12912.

Lee, Woo Yong, Hye Kyung Hong, Soo Kyung Ham, Chang In Kim, and Yong Beom Cho. 2014. “Comparison of Colorectal Cancer in Differentially Established Liver Metastasis Models.” Anticancer Res. 34 (7): 3321–3328. https://www.ncbi.nlm.nih.gov/pubmed/24982336 http://ar.iiarjournals.org/cgi/pmidlookup?view=long&pmid=24982336.

Lupien, Mathieu, Jé Eeckhoute, Clifford A. Meyer, Qianben Wang, Yong Zhang, Wei Li, Jason S. Carroll, X. Shirley Liu, and Myles Brown. 2008. “FoxA1 Translates Epigenetic Signatures into Enhancer-Driven Lineage-Specific Transcription.” Cell 132 (6): 958–970. doi:10.1016/j.cell.2008.01.018. http://www.cell.com/abstract/S0092-8674(08)00118-9 http://dx.doi.org/10.1016/j.cell.2008.01.018 https://www.ncbi.nlm.nih.gov/pubmed/18358809.

Makova, Kateryna D. and Ross C. Hardison. 2015. “The Effects of Chromatin Organization on Variation in Mutation Rates in the Genome; PMC4500049.” Nat.Rev.Genet. 16 (4): 213–223. doi:10.1038/nrg3890. http://dx.doi.org/10.1038/nrg3890 https://www.ncbi.nlm.nih.gov/pubmed/25732611 https://www.ncbi.nlm.nih.gov/pmc/articles/PMC4500049.

Mansour, Marc R., Brian J. Abraham, Lars Anders, Alla Berezovskaya, Alejandro Gutierrez, Adam D. Durbin, Julia Etchin, et al. 2014. “Oncogene Regulation. an Oncogenic Super-Enhancer Formed through Somatic Mutation of a Noncoding Intergenic Element; PMC4720521.” Science 346 (6215): 1373–1377. doi:10.1126/science.1259037. http://dx.doi.org/10.1126/science.1259037 https://www.ncbi.nlm.nih.gov/pubmed/25394790 https://www.ncbi.nlm.nih.gov/pmc/articles/PMC4720521 http://www.sciencemag.org/cgi/pmidlookup?view=long&pmid=25394790 https://www.ncbi.nlm.nih.gov/pmc/articles/PMC4720521/.

Markowitz, S., J. Wang, L. Myeroff, R. Parsons, L. Sun, J. Lutterbaugh, R. S. Fan, E. Zborowska, K. W. Kinzler, and B. Vogelstein. 1995. “Inactivation of the Type II TGF-Beta Receptor in Colon Cancer Cells with Microsatellite Instability.” Science 268 (5215): 1336–1338. https://www.ncbi.nlm.nih.gov/pubmed/7761852 http://www.sciencemag.org/cgi/pmidlookup?view=long&pmid=7761852.

Nakanishi, Yoshito, Kimberly Walter, Jill M. Spoerke, Carol O’Brien, Ling Y. Huw, Garret M. Hampton, and Mark R. Lackner. 2016. “Activating Mutations in PIK3CB Confer Resistance to PI3K Inhibition and Define a Novel Oncogenic Role for p110ß.” Cancer Res. 76 (5): 1193–1203. doi:10.1158/0008-5472.CAN-15-2201. http://dx.doi.org/10.1158/0008-5472.CAN-15-2201 https://www.ncbi.nlm.nih.gov/pubmed/26759240 http://cancerres.aacrjournals.org/cgi/pmidlookup?view=long&pmid=26759240.

Polak, Paz, Rosa Karlic, Amnon Koren, Robert Thurman, Richard Sandstrom, Michael Lawrence, Alex Reynolds, et al. 2015. “Cell-of-Origin Chromatin Organization Shapes the Mutational Landscape of Cancer; PMC4405175.” Nature 518 (7539): 360–364. doi:10.1038/nature14221. http://dx.doi.org/10.1038/nature14221 https://www.ncbi.nlm.nih.gov/pubmed/25693567 https://www.ncbi.nlm.nih.gov/pmc/articles/PMC4405175 https://www.ncbi.nlm.nih.gov/pmc/articles/PMC4405175/.

Pomerantz, Mark M., Fugen Li, David Y. Takeda, Romina Lenci, Apurva Chonkar, Matthew Chabot, Paloma Cejas, et al. 2015. “The Androgen Receptor Cistrome is Extensively Reprogrammed in Human Prostate Tumorigenesis; PMC4707683.” Nat.Genet. 47 (11): 1346–1351. doi:10.1038/ng.3419. http://dx.doi.org/10.1038/ng.3419 https://www.ncbi.nlm.nih.gov/pubmed/26457646 https://www.ncbi.nlm.nih.gov/pmc/articles/PMC4707683.

Puente, Xose S., Silvia Beà, Rafael Valdés-Mas, Neus Villamor, Jesús Gutiérrez-Abril, José Martín-Subero I., Marta Munar, et al. 2015. “Non-Coding Recurrent Mutations in Chronic Lymphocytic Leukaemia.” Nature 526 (7574): 519–524. doi:10.1038/nature14666. http://dx.doi.org/10.1038/nature14666https://www.ncbi.nlm.nih.gov/pubmed/26200345.

Roe, Jae-Seok, Chang-Il Hwang, Tim D. D. Somerville, Joseph P. Milazzo, Eun Jung Lee, Brandon Da Silva, Laura Maiorino, et al. 2017. “Enhancer Reprogramming Promotes Pancreatic Cancer Metastasis; PMC5726277.” Cell 170 (5): 888.e20. doi:10.1016/j.cell.2017.07.007. http://dx.doi.org/10.1016/j.cell.2017.07.007 https://www.ncbi.nlm.nih.gov/pubmed/28757253 https://www.ncbi.nlm.nih.gov/pmc/articles/PMC5726277 https://linkinghub.elsevier.com/retrieve/pii/S0092-8674(17)30814-0.

Santarius, Thomas, Janet Shipley, Daniel Brewer, Michael R. Stratton, and Colin S. Cooper. 2010. “A Census of Amplified and Overexpressed Human Cancer Genes.” Nat.Rev.Cancer 10 (1): 59–64. doi:10.1038/nrc2771. http://dx.doi.org/10.1038/nrc2771 https://www.ncbi.nlm.nih.gov/pubmed/20029424.

Satoh, Kennichi, Shin Hamada, Kenji Kimura, Atsushi Kanno, Morihisa Hirota, Jun Umino, Wataru Fujibuchi, et al. 2008. “Up-Regulation of MSX2 Enhances the Malignant Phenotype and is Associated with Twist 1 Expression in Human Pancreatic Cancer Cells; PMC2276419.” Am.J.Pathol. 172 (4): 926–939. doi:10.2353/ajpath.2008.070346. http://dx.doi.org/10.2353/ajpath.2008.070346 https://www.ncbi.nlm.nih.gov/pubmed/18349132 https://www.ncbi.nlm.nih.gov/pmc/articles/PMC2276419 https://linkinghub.elsevier.com/retrieve/pii/S0002-9440(10)61855-X https://www.ncbi.nlm.nih.gov/pmc/articles/PMC2276419/.

Schuster-Böckler, Benjamin and Ben Lehner. 2012. “Chromatin Organization is a Major Influence on Regional Mutation Rates in Human Cancer Cells.” Nature 488: 504. doi:10.1038/nature11273. http://dx.doi.org/10.1038/nature11273 https://www.nature.com/articles/nature11273.

Sebastian, Alvaro and Bruno Contreras-Moreira. 2014. “footprintDB: A Database of Transcription Factors with Annotated Cis Elements and Binding Interfaces.” Bioinformatics 30 (2): 258–265. doi:10.1093/bioinformatics/btt663. http://dx.doi.org/10.1093/bioinformatics/btt663https://www.ncbi.nlm.nih.gov/pubmed/24234003 https://academic.oup.com/bioinformatics/article-lookup/doi/10.1093/bioinformatics/btt663.

Supek, Fran and Ben Lehner. 2015. “Differential DNA Mismatch Repair Underlies Mutation Rate Variation Across the Human Genome; PMC4425546.” Nature521 (7550): 81–84. doi:10.1038/nature14173. http://dx.doi.org/10.1038/nature14173 https://www.ncbi.nlm.nih.gov/pubmed/25707793 https://www.ncbi.nlm.nih.gov/pmc/articles/PMC4425546 https://www.ncbi.nlm.nih.gov/pmc/articles/PMC4425546/.

Yan, Jian, Martin Enge, Thomas Whitington, Kashyap Dave, Jianping Liu, Inderpreet Sur, Bernhard Schmierer, et al. 2013. “Transcription Factor Binding in Human Cells Occurs in Dense Clusters Formed Around Cohesin Anchor Sites.” Cell 154 (4): 801–813. http://www.sciencedirect.com/science/article/pii/S0092867413009422.

Zhang, Xiaoyang, Peter S. Choi, Joshua M. Francis, Marcin Imielinski, Hideo Watanabe, Andrew D. Cherniack, and Matthew Meyerson. 2016. “Identification of Focally Amplified Lineage-Specific Super-Enhancers in Human Epithelial Cancers; PMC4857881.” Nat.Genet. 48 (2): 176–182. doi:10.1038/ng.3470. http://dx.doi.org/10.1038/ng.3470. https://www.ncbi.nlm.nih.gov/pubmed/26656844 https://www.ncbi.nlm.nih.gov/pmc/articles/PMC4857881.

